# Examining clustered somatic mutations with SigProfilerClusters

**DOI:** 10.1101/2022.02.11.480117

**Authors:** Erik N. Bergstrom, Mousumy Kundu, Noura Tbeileh, Ludmil B. Alexandrov

## Abstract

**Summary:** Clustered mutations are found in the human germline as well as in the genomes of cancer and normal somatic cells. Clustered events can be imprinted by a multitude of mutational processes, and they have been implicated in both cancer evolution and development disorders. Existing tools for identifying clustered mutations have been optimized for a particular subtype of clustered event and, in most cases, relied on a predefined inter-mutational distance (IMD) cutoff combined with a piecewise linear regression analysis. Here we present SigProfilerClusters, an automated tool for detecting all types of clustered mutations by calculating a sample-dependent IMD threshold using a simulated background model that takes into account extended sequence context, transcriptional strand asymmetries, and regional mutation densities. SigProfilerClusters disentangles all types of clustered events from non-clustered mutations and annotates each clustered event into an established subclass, including the widely used classes of doublet-base substitutions, multi-base substitutions, *omikli*, and *kataegis*. SigProfilerClusters outputs non-clustered mutations and clustered events using standard data formats as well as provides multiple visualizations for exploring the distributions and patterns of clustered mutations across the genome.

**Availability:** SigProfilerClusters is freely available at https://github.com/AlexandrovLab/SigProfilerClusters with support across most operating systems and extensive documentation at https://osf.io/qpmzw/wiki/home/.

## INTRODUCTION

Mutations are found on the genomes of all cells in the human body (Martincorena and Campbell, 2015; Stratton, et al., 2009). Most single-base substitutions and small insertions and deletions (indels) accumulate independently across the genome, but a subset of mutations cluster in a non-random manner (Lawrence, et al., 2013; Supek and Lehner, 2017). Previous studies have revealed that clustered mutations are imprinted by a plethora of endogenous and exogenous mutational processes (Alexandrov, et al., 2020; Boichard, et al., 2017; Brash, 2015; Buisson, et al., 2019; Chan, et al., 2015; Chen, et al., 2013; Consortium, 2020; Mas-Ponte and Supek, 2020; Matsuda, et al., 1998; Nik-Zainal, et al., 2012; Nik-Zainal, et al., 2019; Pfeifer, et al., 2005; Roberts, et al., 2013; Roberts, et al., 2012; Supek and Lehner, 2017; Taylor, et al., 2013; Wang, et al., 2020). Some clustered mutations have been implicated in cancer evolution (Bergstrom, et al., 2022; Chen, et al., 2013; Consortium, 2020; Mas-Ponte and Supek, 2020; Supek and Lehner, 2017; Taylor, et al., 2013), while *de novo* clustered mutations have been identified in the human germline and shown to contribute to developmental disorders (Kaplanis, et al., 2019; Veltman and Brunner, 2012). In recent years, sets of simultaneously occurring clustered substitutions have been further subclassified into independent events (Bergstrom, et al., 2022; Mas-Ponte and Supek, 2020), including: *(i)* doublet-base substitutions (DBSs); *(ii)* multi-base substitutions (MBSs); *(iii)* diffuse hypermutation termed *omikli*; *(iv)* longer strand-coordinated events termed *kataegis*; and *(v)* recurrent hypermutation of extra-chromosomal DNA (ecDNA) termed *kyklonas*.

Traditional methods separate clustered mutations based upon a predefined inter-mutational distance (IMD) threshold typically between 1 and 2 kilobases (Alexandrov, et al., 2020; Alexandrov, et al., 2013; Chan, et al., 2015; D’Antonio, et al., 2016; Maciejowski, et al., 2020; Nik-Zainal, et al., 2019; Taylor, et al., 2013). Many of these approaches utilize a piece-wise linear regression to segment each chromosome, which, in most cases, is optimized for calling larger strand-coordinated *kataegic* events (**Supplementary Fig. 1**) (Alexandrov, et al., 2013; Lin, et al., 2021; Yin, et al., 2020). Most existing methods have also ignored confounding effects attributed to localized differences in mutation rates, copy number alterations, or the mutational burden across each chromosome within a given sample leading to an accumulation of false positive clustered events (**Supplementary Fig. 1**). Further, the majority of existing tools focus on detecting only a specific class of clustered events including doublet-base substitutions and multi-nucleotide variants (Chen, et al., 2013; Matsuda, et al., 1998; Wang, et al., 2020), *kataegis* (D’Antonio, et al., 2016; Lin, et al., 2021; Taylor, et al., 2013), or APOBEC3-associated events (Chan, et al., 2015; Nik-Zainal, et al., 2012) while ignoring the larger landscape of clustered mutations. For example, a recent study (Mas-Ponte and Supek, 2020) developed an algorithm focused on the detection of APOBEC3-associated *omikli* and *kataegis* events in cancer genomes by incorporating simulations of somatic mutations and estimates of cancer cell fractions.

Separation and classification of clustered events is required to fully elucidate the mutational processes operating in cancer and normal somatic cells (Bergstrom, et al., 2022; Supek and Lehner, 2017). Here we present SigProfilerClusters, a tool to comprehensively characterize and subclassify clustered mutations from the complete catalog of mutations within the genome of a single sample (**Fig. 1*a***). SigProfilerClusters classifies all types of clustered mutations, including *(i)* doublet-base substitutions; *(ii)* multi-base substitutions; *(iii) omikli*; *(iv) kataegis*; and *(v)* clustered small insertions and deletions (indels). The tool calculates a sample-dependent IMD threshold that considers regional differences in mutation rates, variant allele fractions and cancer cell fractions of adjacent mutations to reduce the false positive rate and provides visualizations for downstream analyses (**Fig. 1*b&c***; **Supplementary Fig. 1**). Further, SigProfilerClusters integrates within the larger suite of SigProfiler tools (Bergstrom, et al., 2020; Bergstrom, et al., 2019; Islam, et al., 2020) to facilitate downstream mutational signature analysis of both non-clustered and clustered single-base substitutions and indels, thus, allowing the accurate detection of mutational processes giving rise to even low levels of clustered events (**Fig. 1*d***) (Bergstrom, et al., 2019; Bergstrom, et al., 2022; Islam, et al., 2020).

## METHODS

SigProfilerClusters derives an IMD cutoff that is unlikely to occur purely by chance given the observed mutational burden and the mutational patterns within the genome of a given sample. To calculate the genome-dependent IMD, the tool leverages SigProfilerSimulator (Bergstrom, et al., 2020) to generate background models by randomizing the distribution of mutations across the genome. By default, the genome of each sample is simulated 100 times in order to derive 95% confidence intervals for the expected genomic mutational landscape, with every simulation maintaining the penta-nucleotide sequence context for each substitution, the ratio of all mutations in genic and inter-genic regions, the transcriptional strand asymmetries of all mutations in genic regions, and the mutational burden on each chromosome (Bergstrom, et al., 2020; Bergstrom, et al., 2019). Importantly, this randomization procedure is highly customizable (Bergstrom, et al., 2020) and can be altered based upon the needs of a given study design, thus, allowing the incorporation of other factors that affect the accumulation of mutations such as nucleosome occupancy, presence of histone modifications, and many others. A binary search algorithm is implemented to efficiently derive the global IMD threshold for each genome. The final global IMD threshold is selected by ensuring that 90% of mutations below the chosen cutoff are unlikely to appear by chance given the simulated distribution of mutations (q-value<0.01; **Supplementary Fig. 1**) with a maximum global IMD cutoff of 10 kilobases. The algorithm also considers regional heterogeneities of mutation rates, generally associated with replication timing (Stamatoyannopoulos, et al., 2009) or differential gene expression (Buisson, et al., 2019; Hess, et al., 2019; Lawrence, et al., 2013; Pleasance, et al., 2010; Polak, et al., 2015), by correcting for variance in clonality as well as variance in both mutation-dense and mutation-poor regions using a sliding genomic window (default size of 1 megabase). Specifically, an additional regional IMD cutoff is corrected within each genomic window based upon the fold difference between the number of real and the number of simulated mutations, while maintaining the original criteria of less than 10% of mutations below the IMD cutoff appearing by chance (q-alue<0.01). Lastly, when data are available, SigProfilerClusters ensures that adjacent mutations are in the same cells by introducing a maximum difference in variant allele frequencies (VAF) or cancer cell fraction (CCF), which incorporates copy-number changes, below a certain threshold (default cutoff value of 0.10 and 0.25; respectively).

After identifying the set of clustered mutations, SigProfilerClusters subclassifies each clustered substitution into a single category of previously established clustered events (Bergstrom, et al., 2022; Mas-Ponte and Supek, 2020). Briefly, all clustered substitutions with consistent VAFs or consistent CCFs are classified into one of four categories. Two mutations with an IMD of 1 are classified as *doublet-base substitutions*, while clusters of three or more adjacent mutations each with an IMD of 1 are classified as *multi-base substitutions*. Clusters of two or three mutations with IMDs less than the sample-dependent cutoff and with at least a single IMD greater than 1 are classified as *omikli* (Bergstrom, et al., 2022), while clusters of four or more mutations with IMDs less than the sample-dependent cutoff and with at least a single IMD greater than 1 are classified as *kataegis* (Bergstrom, et al., 2022). All remaining clustered mutations with inconsistent VAFs or CCFs are classified as *other*. Clustered indels are not subclassified into different categories due to a lack of previously defined subtypes.

## USAGE

SigProfilerClusters is freely available as a Python package, distributed under the permissive BSD-2 clause license, and can be used on most operating systems including Windows, MacOS, and Linux-based machines. The tool is compatible with large-scale deployments on high-performance computing clusters as well as on cloud infrastructures such as Amazon Web Services (AWS). Input data can be provided in the form of common mutation formats including the Variant Call Format (VCF), the Mutation Annotation Format (MAF), or in the form of a simple text file. The output of SigProfilerClusters results in the partitioning of all mutations into a clustered or non-clustered directory. All clustered mutations are then classified into distinct subcategories of events and provided individually in VCF files for downstream visualization and analyses. The output for each subclass of clustered event can be directly utilized by additional SigProfiler tools including SigProfilerExtractor for mutational signature analysis (Islam, et al., 2020) and SigProfilerPlotting for examining patterns of somatic mutations (Bergstrom, et al., 2019). The results for each sample are also summarized using two individual visualizations that include: *(i)* a rainfall plot depicting the minimum global IMD between all adjacent mutations, where each individual set of adjacent mutations is colored based upon its clustered classification; and *(ii)* a multi-panel figure that displays the mutational patterns across all mutations, clustered mutations, and non-clustered mutations, separately along with the distribution of IMDs across the real and simulated data for each sample (**Fig. 1*a***).

## CONCLUSION

Elucidating the compendium of clustered somatic mutations in the genome of a sample allows further understanding of the mutational process that give rise to these events and can provide novel insights into disease etiology (Bergstrom, et al., 2022; Mas-Ponte and Supek, 2020; Supek and Lehner, 2017). Previous studies have traditionally interrogated the complete mutational catalogs of cancer genomes, which can lead to the inability to detect processes active at low levels or those which have been transiently activated. Our prior analysis of clustered mutations (Bergstrom, et al., 2022) have revealed an enrichment of clustered mutations within known cancer driver events, hypermutation of extrachromosomal DNA fueling the evolution of cancers, and ultimately, resulting in a differential patient outcome. Here we provide SigProfilerClusters, an automated and freely available Python based tool that comprehensively identifies and classifies clustered mutations enabling users to interrogate the mutational processes giving rise to such events.

**Figure 1.**
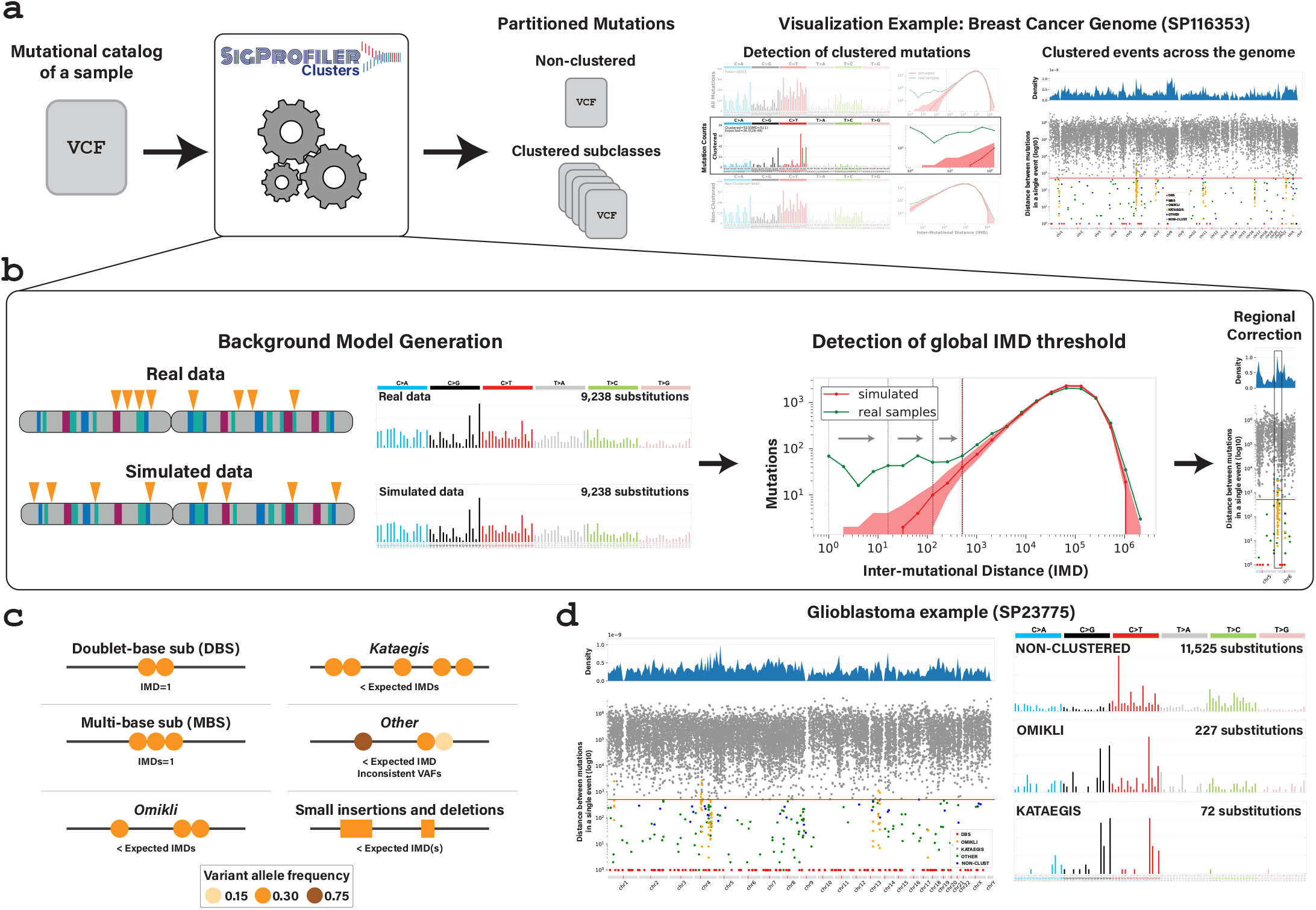
Detection and characterization of clustered mutations with SigProfilerClusters. ***a)*** An example workflow used to detect clustered mutations in a single cancer genome. As an input, SigProfilerClusters accepts common formats for mutations, such as ones in the variant calling format (VCF), and the tool separates all clustered mutations from the complete mutational catalog of the provided sample. Final partitions of mutations in the sample are outputted as VCF files and visualized using the mutational spectra of all mutations, only clustered mutations, and only non-clustered mutations along with a rainfall plot commonly used to show the distribution of inter-mutational distances across a cancer genome (Alexandrov, et al., 2013; Bergstrom, et al., 2022; Nik-Zainal, et al., 2012). ***b)*** Schematic demonstrating the process of calculating a sample dependent IMD threshold to separate clustered from non-clustered mutations across each genome. A binary search algorithm is used to efficiently detect the optimal global IMD threshold for each sample. Detection of the global IMD threshold is illustrated using grey arrows. Regional corrections are performed to identify local IMD thresholds based on variance of mutation rates across the genome. ***c)*** Every clustered mutation is classified into a single subcategory of clustered event. ***d)*** Rainfall plot illustrating the distribution of IMDs across a single glioblastoma sample (*left*). The mutational spectra for *omikli* and *kataegic* events reveal a different mutational pattern compared to the pattern of all non-clustered somatic mutations (*right*).

## SUPPLEMENTARY FIGURE LEGENDS

**Supplementary Figure 1. Benchmarking of existing tools for detecting clustered mutations.**

***a***) Assessing the degree of overlap between two tools that detect clustered mutations and SigProfilerCusters. P-MACD locates regions of clustered mutations by implementing a negative binomial distribution to model the probability of observing a given cluster within a given window of the genome. When calculating p-values, this model assumes that all mutations occur randomly across the genome limiting clustered events to those that occur at most 10 kilobases (kb) from one another. Kataegis implement a piece-wise linear model with set thresholds requiring all clustered events to be composed of 6 or more mutations with an average intermutational distance of 1 kb between adjacent mutations. All tools were applied to 2,703 whole-genome sequenced samples from PCAWG and were split into low (TMB<1), intermediate (1<=TMB<10), and high (TMB>=10) TMB. ***b)*** The percentage of mutations overlapping each subclass of clustered events called by SigProfilerClusters (*left*), the percentage of mutations that were missed by each of the two tools that were called by SigProfilerClusters (*middle*), and the percentage of total mutations called by each tool that were missed by SigProfilerClusters separated by low, intermediate, and high TMB (*right*). ***c)*** Schematic workflow to assess the false positive rate of each tool using simulations of 211 breast cancer genomes from PCAWG. ***d)*** The average number of false positive mutations per simulated sample detected across the simulated breast cancer dataset from PCAWG. Kataegis did not detect any *kataegic* clustered events within the simulated data using the tools’ default parameters. Similarly, SigProfilerClusters did not detect any *kataegic* clustered events within these simulated data.

## Contributions

ENB developed the Python code and wrote the manuscript. MK performed all benchmarking. ENB, MK, and NT tested and documented the code. LBA supervised the overall development of the code, benchmarking, and writing of the manuscript. All authors read and approved the final manuscript.

