## Supplementary figures and images for "Examining clustered somatic mutations with SigProfilerClusters"

### Supplementary Figure 1

#### Supplementary Figure 1. Benchmarking of existing tools that detect clustered mutations

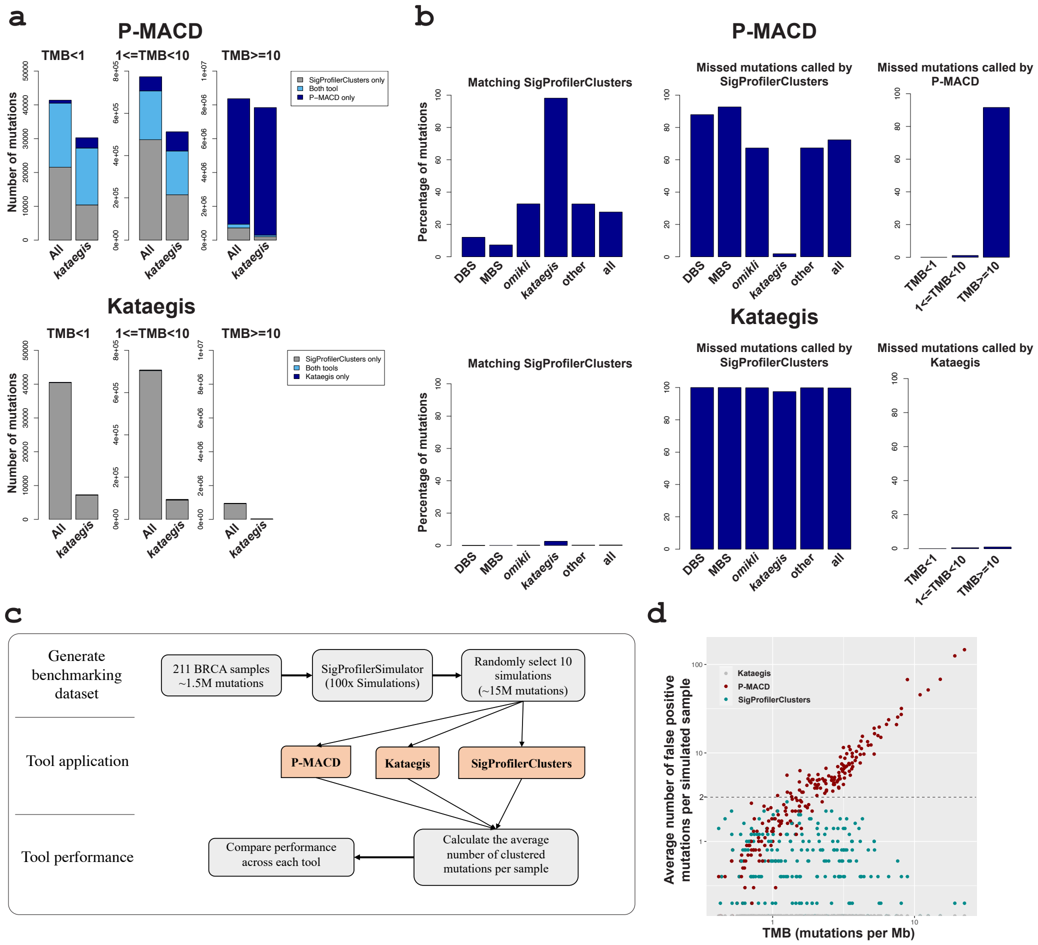
